# Genome-wide association studies on endometriosis and endometriosis-related infertility

**DOI:** 10.1101/401448

**Authors:** Geneviève Galarneau, Pierre Fontanillas, the Celmatix Research Team, the 23andMe Research Team, Caterina Clementi, Tina Hu-Seliger, David-Emlyn Parfitt, Joyce Y. Tung, Piraye Yurttas Beim

## Abstract

Endometriosis affects ∼10% of women of reproductive age. It is characterized by the growth of endometrial-like tissue outside the uterus and is frequently associated with severe pain and infertility. We performed the largest endometriosis genome-wide association study (GWAS) to date, with 37,183 cases and 251,258 controls. All women were of European ancestry. We also performed the first GWAS of endometriosis-related infertility, including 2,969 cases and 3,770 controls. Our endometriosis GWAS study replicated, at genome-wide significance, seven loci identified in previous endometriosis GWASs (*CELA3A-CDC42, SYNE1, KDR, FSHB-ARL14EP, GREB1, ID4*, and *CEP112*) and identified seven new candidate loci with genome-wide significance (*NGF, ATP1B1-F5, CD109, HEY2, OSR2-VPS13B, WT1*, and *TEX11-SLC7A3*). No loci demonstrated genome-wide significance for endometriosis-related infertility, however, the three most strongly associated loci (*MCTP1, EPS8L3-CSF1*, and *LPIN1*) were in or near genes associated with female fertility or embryonic lethality in model organisms. These results reveal new candidate genes with potential involvement in the pathophysiology of endometriosis and endometriosis-related infertility.

## Introduction

Endometriosis affects ∼10% of women of reproductive age and is characterized by the growth of endometrial-like tissue outside the uterus. The ovary is the organ that is most often affected by endometriotic lesions [1], but other organs, including the bowel, fallopian tubes, or bladder, can be affected. Endometriosis is an estrogen-dependent inflammatory disease, with the endometriotic lesions continuing to respond to estrogen and to thicken following the menstrual cycle, creating inflammation and scar tissue called adhesions. The adhesions and associated inflammation are thought to underlie the symptoms of endometriosis, which include pelvic pain, dysmenorrhea, and pain during intercourse.

Endometriosis is also frequently associated (30-50%) with infertility [2]. Women with endometriosis have a lower monthly fecundity rate than fertile controls [3]. A meta-analysis of 22 published studies showed that women with endometriosis undergoing in vitro fertilization (IVF) had significantly reduced oocyte retrieval, fertilization, and implantation rates, as well as a significantly lower chance of achieving pregnancy, compared to patients with tubal factor [4]. Women with endometriosis-related infertility tend to have a longer time to natural conception than women with idiopathic infertility [5, 6] and longer time to conception through artificial insemination than women with no female infertility factors [7]. Even mild endometriosis can affect a woman’s fertility, and infertility-related endometriosis may be associated with endometriosis grade—for example, pregnancy rates following IVF were significantly lower in women with severe endometriosis than in women with mild endometriosis [4]. Endometriosis may also reduce oocyte quality, as suggested by a study in which women who received oocytes donated by women with endometriotic ovaries had a lower implantation rate than those receiving oocytes donated by women without known endometriosis [8].

The causal mechanisms of endometriosis-related infertility are not fully understood. Different aspects of endometriosis may lead to infertility, and the exact cause of endometriosis-related infertility in each patient may depend on the course of her specific pathology. Proposed mechanisms for endometriosis-related infertility include ovarian-tubal dysfunction, immunological disorder, abnormal peritoneal environment, and dysregulated endometrial function [9]. Ovarian-tubal dysfunction could be caused by a distortion of the ovary and/or fallopian tube affected by endometriotic lesions, ovulation failure, or abnormal follicle development. An immunological disorder could be caused in part by the IgG and IgM anti-endometrial antibodies, which have been detected in 60% of endometriosis patients and could impair implantation [10]. Abnormalities in the peritoneal environment could include increased peritoneal fluids and a higher concentration of cytokines and/or activated macrophages. Women with endometriosis-related infertility have an increased volume of peritoneal fluid, and this fluid may have increased levels of inflammatory cytokines due to the activation of macrophages by the endometriotic lesions. Activated macrophages increase reactive oxygen species in the peritoneal fluid of women with endometriosis [11], causing oxidative stress, which has been associated with negative outcomes in assisted reproductive technology fertilization [12]. For example, this increase in cytokines may reduce oocyte quality, sperm motility, and tubal motility, and impair embryo development. Finally, endometrial receptivity is a crucial process to achieve pregnancy and involves complex regulation of hormones, cytokines, and adhesion molecules. This process may be dysregulated in women with endometriosis.

Laparoscopy is currently the only way to diagnose endometriosis with certainty. It involves a surgical procedure in which a surgeon visualizes the endometriotic lesions or collects tissue samples for histologic assessment. Laparoscopy is often used to excise the endometriotic lesions to relieve pain, but symptoms often return in following years. Because endometriosis pain levels and disease severity are not directly correlated, the diagnosis and grading of endometriosis without surgery is difficult. However, because of the invasive nature of the diagnostic procedure, women with suspected endometriosis are often treated with hormone therapy without a definitive diagnosis. Despite its high prevalence and its association with severe pain and infertility, treatment for endometriosis remains limited. Surgical and medical management of endometriosis can help treat symptoms. Surgical management increases the monthly fecundity rate of women with endometriosis but even with surgical management, it remains much lower than in the general population [13]. Hormone treatments include oral contraceptives, androgenic agents, progestins, and gonadotropin-releasing hormone analogs, but they are not always effective.

Endometriosis is a complex trait, with both genetic and environmental factors contributing to the pathophysiology of the disease. The heritability of endometriosis is estimated to be 47-51% [14,15], but the 13 loci identified by previous genome-wide association studies (GWASs) only explain 1.75% of the phenotypic variance [16-23]. Further, few studies have investigated whether genetic factors might be involved in the pathophysiology of endometriosis-related infertility. The identification of novel genetic factors implicated in endometriosis or endometriosis-related infertility could help to elucidate the pathophysiology of endometriosis and to identify potential new drug targets. Here, we conducted a GWAS on endometriosis including 37,183 cases and 251,258 controls and the first GWAS on endometriosis-related infertility, including 2,969 women with endometriosis-related infertility and 3,770 women with endometriosis who reported a first time-to-conception ≤ 6months.

## Materials and Methods

### Participants

Participants in this study were customers of the consumer genetics company 23andMe, Inc., who provided informed consent to participate in research by completing surveys online under a protocol approved by Ethical & Independent Review Services, an independent institutional review board (http://www.eandireview.com) accredited by the Association for the Accreditation of Human Research Protection Programs, Inc. This study was restricted to unrelated female participants with >97% European ancestry. Percentage ancestry was assessed through a local ancestry analysis.

### Genotyping

DNA extraction and genotyping were performed on saliva samples by the National Genetics Institute. Samples were genotyped on one of four custom genome-wide genotyping arrays targeting 560-950k SNPs. Only samples that achieved an average genotyping rate ≥98.5% were included in the analysis.

### Quality control

Genotyped SNPs were excluded if they had a genotyping rate of <90%, did not pass a parent-offspring concordance test, were monomorphic, were not in Hardy-Weinberg equilibrium (p<1×10^-20^), or showed batch effects.

### Imputation

Imputation was performed using the September 2013 release of the 1000 Genomes Project Phase I as reference haplotypes. The phasing was performed with ShapeIt2 and the imputation with Minimac2. Imputed SNPs were flagged and excluded if they had an average r^2^ < 0.5 across genotyping platform or a minimum r^2^ < 0.3 or if they showed strong evidence of imputation batch effect (p<10^-50^).

### Principal components calculations

The principal component analysis was performed using ∼65,000 high quality genotyped variants present in all five genotyping platforms with 1 million of participants from European ancestry randomly sampled across all the genotyping platforms. PC scores for participants not included in the analysis were obtained by projection, combining the eigenvectors of the analysis and the SNP weights.

### Phenotype definition

Case-control ascertainment was based on participants’ survey responses. For the endometriosis case-control GWAS, cases were women who reported being treated for and/or diagnosed with endometriosis. Controls were women who reported not being treated for or diagnosed with endometriosis.

For the endometriosis-related infertility GWAS, both cases and controls were women who reported being treated for and/or diagnosed with endometriosis and who had answered at least one question on time to first conception. The time to first conception was ascertained based on their answers to the question(s): “For how long did you try to conceive? If you are currently trying to conceive, please state how long you have been trying up to this point”, “The first time you tried to conceive a child, for how long were you trying to conceive? If you are currently trying to conceive for the first time, please answer for how long you have been trying up to this point.” The questions regarding time to first conception had multiple choice answers with the following options: 0-6 months, 7-12 months, 13-24 months, 24 months, 24+ months, not successful, and not sure. ‘Infertile’ cases were defined as women who responded with 13-24 months, 24 months, 24+ months, or not successful (35.7% of women with endometriosis in 23andMe database). Women who responded with 7-12 months (13.1%) or I’m not sure were excluded from the analysis (<0.1%). Controls were women who indicated 0-6 months (51.2%) and reported having at least one biological child.

### Statistical analysis

Assuming an additive model, we performed logistic regressions on endometriosis case-control status and on endometriosis-related infertility case-control status with current age, the top five principal components, and the genotyping platform as covariates. The association results were adjusted for the observed genomic control inflation factor in the distribution of the p-values (λ=1.141 and λ=1.011, respectively, Supplementary Figures 1-2).

**Figure 1.**
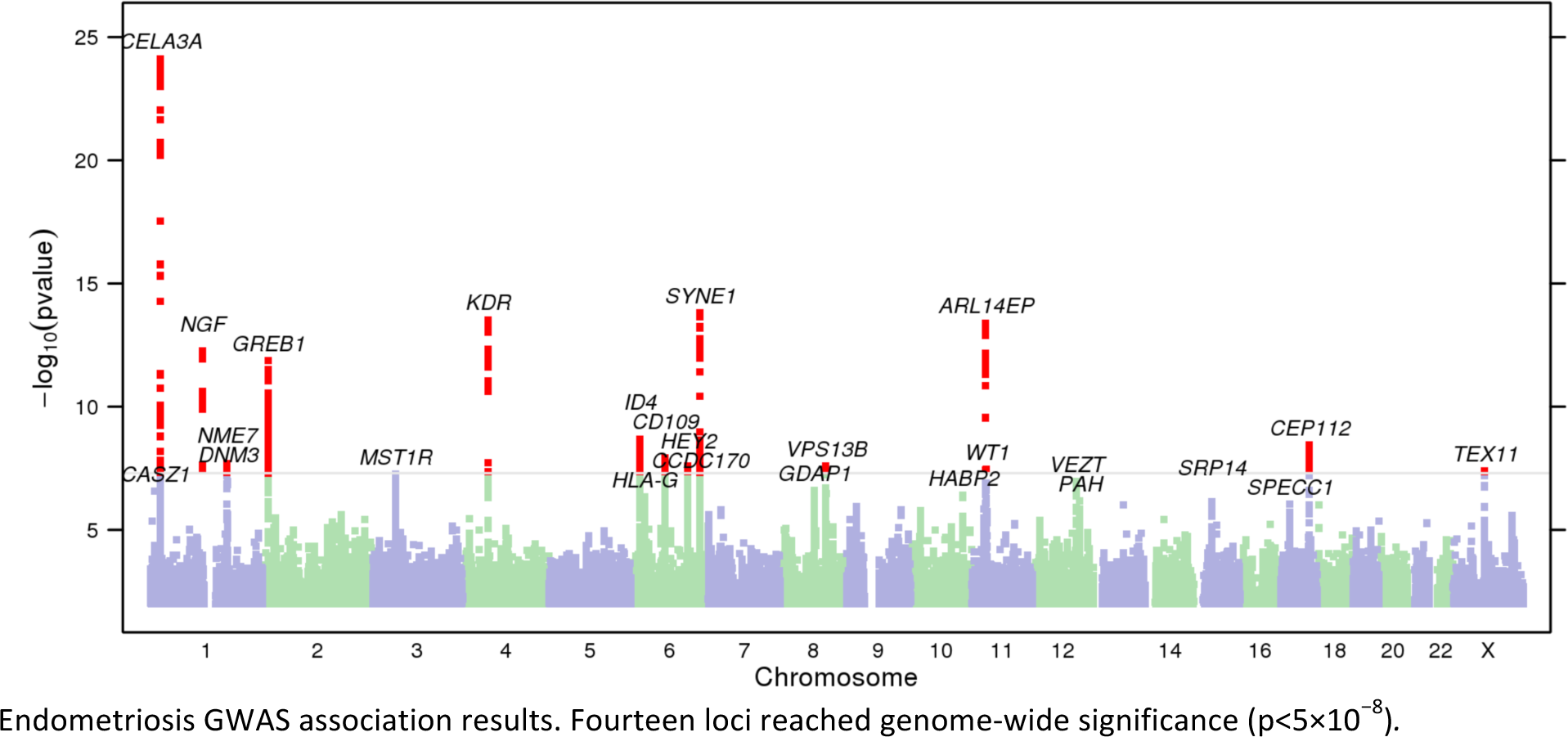
Manhattan plot of endometriosis GWAS.

### Prioritization of putative functional variants

The 99% credible sets were generated by first calculating an approximate Bayes factor (ABF) for each SNP, using the method proposed by Wakefield [24] and assuming a prior variance (W) of 0.1. The credible sets were then estimated from the ABFs, using the method of Maller *et al* [25]. We prioritized genetic variants in the credible sets for further follow-up in GTEx Portal and RegulomeDB and used LDlink [26] to identify potential genetic variants that would not have passed genotyping/imputation QC but would be in high linkage disequilibrium (LD) (r^2^ > 0.8 in individuals of European ancestry in the 1000 Genomes Project).

## Results

Fourteen loci reached genome-wide significance (p<5×10^-8^), including seven loci not reported in previous GWASs (*NGF, ATP1B1-F5, CD109, HEY2, OSR2-VPS13B, WT1*, and *TEX11-SLC7A3*) (Figure 1, Table 3). Among the loci that were previously reported in the literature and did not reach genome-wide significance, four were nominally associated (p<0.05) with a consistent direction of effect (*FGD6-VEZT, RMND1-ESR1, NPVF-NFE2L3*, and *CDKN2B-DMRTA1*) and two had p-values >0.05 (*IGFBP3-TNS3* and *TTC39B*) (Table 4).

**Table 1.**
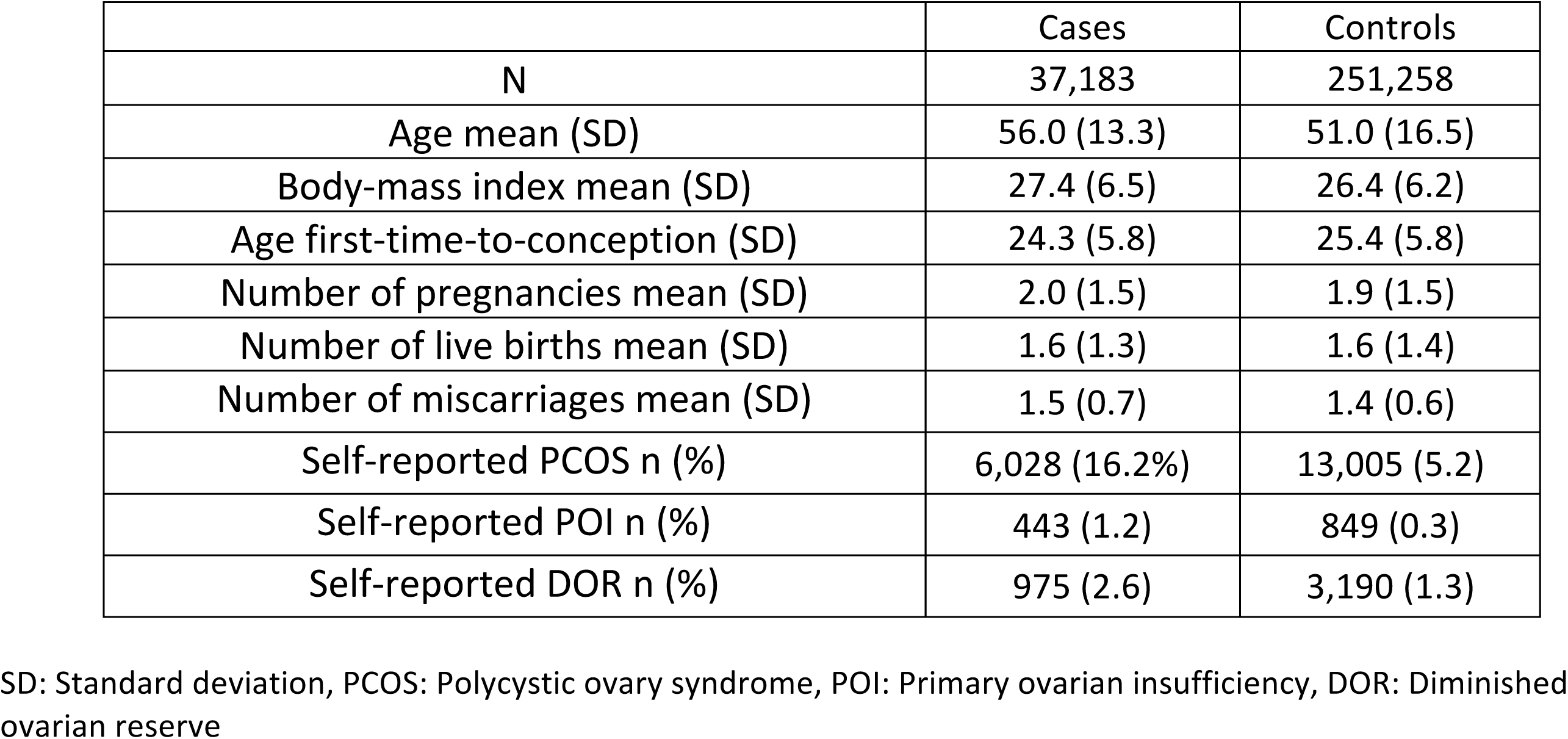
Cohort description for endometriosis case-control GWAS

|  | Cases | Controls |
| --- | --- | --- |
| N | 37,183 | 251,258 |
| Age mean (SD) | 56.0 (13.3) | 51.0 (16.5) |
| Body-mass index mean (SD) | 27.4 (6.5) | 26.4 (6.2) |
| Age first-time-to-conception (SD) | 24.3 (5.8) | 25.4 (5.8) |
| Number of pregnancies mean (SD) | 2.0 (1.5) | 1.9 (1.5) |
| Number of live births mean (SD) | 1.6 (1.3) | 1.6 (1.4) |
| Number of miscarriages mean (SD) | 1.5 (0.7) | 1.4 (0.6) |
| Self-reported PCOS n (%) | 6,028 (16.2%) | 13,005 (5.2) |
| Self-reported POI n (%) | 443 (1.2) | 849 (0.3) |
| Self-reported DOR n (%) | 975 (2.6) | 3,190 (1.3) |

SD: Standard deviation, PCOS: Polycystic ovary syndrome, POI: Primary ovarian insufficiency, DOR: Diminished ovarian reserve

**Table 2.** Cohort description for endometriosis-related infertility GWAS

|  | Cases | Controls |
| --- | --- | --- |
| N | 2,969 | 3,770 |
| Age mean (SD) | 56.0 (13.3) | 51.0 (16.5) |
| Body-mass index mean (SD) | 27.4 (6.5) | 26.4 (6.2) |
| Age first-time-to-conception (SD) | 24.3 (5.8) | 25.4 (5.8) |
| Number of pregnancies mean (SD) | 2.0 (1.5) | 1.9 (1.5) |
| Number of live births mean (SD) | 1.6 (1.3) | 1.6 (1.4) |
| Number of miscarriages mean (SD) | 1.5 (0.7) | 1.4 (0.6) |
| Self-reported PCOS n (%) | 6,028 (16.2%) | 13,005 (5.2) |
| Self-reported POI n (%) | 443 (1.2) | 849 (0.3) |
| Self-reported DOR n (%) | 975 (2.6) | 3,190 (1.3) |

SD: Standard deviation, PCOS: Polycystic ovary syndrome, POI: Primary ovarian insufficiency, DOR: Diminished ovarian reserve

**Table 3.** Endometriosis case-control GWAS genome-wide loci

|  |  |  |  |  |  |  | Endometriosis case-control GWAS |  | Endometriosis-related infertility |  |
| --- | --- | --- | --- | --- | --- | --- | --- | --- | --- | --- |
| Variant id | Chr | Pos (hg19) | Nearest gene(s) | RA | OA | RAF | OR [95% CI] | P | OR [95% CI] | P |
| Novel loci |  |  |  |  |  |  |  |  |  |  |
| rs12030576 | 1 | 115817221 | *NGF* | G | T | 0.65 | 1.07 [1.05-1.09] | $5.2 \times 10^{-13}$ | 1.09 [1.00, 1.18] | 0.03 |
| rs1209731 | 1 | 169324793 | *ATP1B1-F5* | C | T | 0.98 | 1.19 [1.12-1.26] | $2.0 \times 10^{-8}$ | 1.11 [0.86, 1.43] | 0.44 |
| rs1595344 | 6 | 74611632 | *CD109* | G | A | 0.66 | 1.05 [1.03-1.07] | $1.2 \times 10^{-8}$ | 1.04 [0.96, 1.13] | 0.32 |
| rs2226158 | 6 | 125995467 | *HEY2* | G | A | 0.47 | 1.05 [1.03-1.07] | $2.6 \times 10^{-8}$ | 0.95 [0.90, 1.01] | 0.16 |
| rs6468654 | 8 | 100062724 | *OSR2-VPS13B* | C | T | 0.24 | 1.06 [1.04-1.08] | $2.5 \times 10^{-8}$ | 1.00 [0.93, 1.08] | 0.96 |
| rs7924571 | 11 | 32350027 | *WT1* | C | A | 0.77 | 1.06 [1.04-1.08] | $3.5 \times 10^{-8}$ | 0.99 [0.92, 1.08] | 0.90 |
| rs13441059 | X | 70108889 | *TEX11-SLC7A3* | A | G | 0.36 | 1.05 [1.03-1.07] | $4.1 \times 10^{-8}$ | 1.01 [0.93, 1.09] | 0.82 |
| Previously reported loci |  |  |  |  |  |  |  |  |  |  |
| rs80173514 | 1 | 22354538 | *CELA3A-CDC42* | A | C | 0.15 | 1.13 [1.10-1.15] | $7.5 \times 10^{-25}$ | 1.15 [1.05, 1.27] | $2.7 \times 10^{-3}$ |
| rs17215781 | 6 | 152570274 | *SYNE1* | A | G | 0.92 | 1.13 [1.10-1.17] | $1.5 \times 10^{-14}$ | 1.16 [1.01, 1.33] | 0.04 |
| rs9312658 | 4 | 56005526 | *KDR* | C | A | 0.67 | 1.07 [1.05-1.09] | $2.9 \times 10^{-14}$ | 1.09 [1.00, 1.17] | 0.03 |
| rs10616649 | 11 | 30349488 | *FSHB-ARL14EP* | ACTCTA | - | 0.84 | 1.09 [1.07-1.12] | $4.0 \times 10^{-14}$ | 0.94 [0.85, 1.04] | 0.23 |
| rs11674184 | 2 | 11721535 | *GREB1* | T | G | 0.61 | 1.07 [1.05-1.08] | $1.4 \times 10^{-12}$ | 0.99 [0.92, 1.07] | 0.83 |
| rs6456259 | 6 | 19761718 | *ID4* | G | A | 0.16 | 1.07 [1.05-1.10] | $2.1 \times 10^{-9}$ | 0.95 [0.86, 1.04] | 0.25 |
| rs746628 | 17 | 63850547 | *CEP112* | T | C | 0.69 | 1.06 [1.04-1.08] | $3.7 \times 10^{-9}$ | 0.99 [0.91, 1.07] | 0.70 |

Genome-wide significant loci in endometriosis case-control GWAS
Chr: chromosome, Pos: position, RA: risk allele, OA: Other allele, RAF: Risk allele frequency, OR: Odds ratio, P: P-value

**Table 4.** Endometriosis case-control GWAS association results for other previously reported loci

| Variant id | Chr | Pos (hg19) | Nearest gene(s) | RA | OA | RAF | Endometriosis case-control GWAS |  | Endometriosis-related infertility |  |
| --- | --- | --- | --- | --- | --- | --- | --- | --- | --- | --- |
|  |  |  |  |  |  |  | OR [95% CI] | P | OR [95% CI] | P |
| rs10859853 | 12 | 95614953 | *FGD6-VEZT* | C | T | 0.37 | 1.05 [1.03, 1.06] | $1.1 \times 10^{-7}$ | 1.04 [0.96, 1.12] | 0.28 |
| rs4762326 | 12 | 95668951 | | T | C | 0.48 | 1.04 [1.02, 1.06] | $1.6 \times 10^{-6}$ | 1.02 [0.96, 1.08] | 0.49 |
| rs4870024 | 6 | 151830769 | *RMND1-ESR1* | C | T | 0.19 | 1.06 [1.04, 1.08] | $1.8 \times 10^{-7}$ | 0.95 [0.88, 1.03] | 0.24 |
| rs1971256 | 6 | 151816011 | | C | T | 0.21 | 1.05 [1.03, 1.07] | $9.2 \times 10^{-7}$ | 0.96 [0.90, 1.04] | 0.35 |
| rs12700667 | 1 | 25901639 | *NPVF-NFE2L3* | A | G | 0.75 | 1.04 [1.02, 1.06] | $6.3 \times 10^{-5}$ | 1.05 [0.97, 1.13] | 0.24 |
| rs1537377 | 9 | 22169700 | *CDKN2B-DMRTA1* | C | T | 0.38 | 1.04 [1.03, 1.06] | $1.6 \times 10^{-6}$ | 0.97 [0.90, 1.05] | 0.48 |
| rs74491657 | 7 | 46947633 | *IGFBP3-TNS3* | G | A | 0.90 | 1.02 [1.00, 1.04] | 0.12 | 1.00 [0.89, 1.13] | 0.95 |
| rs519664 | 9 | 15246652 | *TTC39B* | T | C | 0.21 | 1.01 [0.99, 1.03] | 0.34 | 0.96 [0.89, 1.04] | 0.35 |

Association results for other previously reported endometriosis loci
Chr: chromosome, Pos: position, RA: risk allele, OA: Other allele, RAF: Risk allele frequency, OR: Odds ratio, P: P-value

No loci reached genome-wide association with endometriosis-related infertility (Figure 2, Table 5). Among the 14 loci that reached genome-wide significance in the endometriosis case-control GWAS, four (*NGF, CELA3A-CDC42, SYNE1*, and *KDR*) reached nominal association (p<0.05) with shared risk alleles for both endometriosis and endometriosis-related infertility (Table 3). Five loci reached suggestive levels of association (p<1×10^-6^) with endometriosis-related infertility (*MCTP1, EPS8L3-CSF1, LPIN1, GRM8*, and *FSTL5-NAF1*) (Table 5).

**Table 5.** Top loci associated with endometriosis-related infertility

| Variant id | Chr | Pos (hg19) | Nearest gene(s) | RA | OA | RAF | OR [95% CI] | P |
| --- | --- | --- | --- | --- | --- | --- | --- | --- |
| rs201312237 | 5 | 94607502 | *MCTP1* | ATAG | - | $8.8 \times 10^{-3}$ | 2.96 [1.93-4.55] | $2.3 \times 10^{-7}$ |
| rs182207890 | 1 | 110356869 | *EPS8L3-CSF1* | C | T | 0.998 | Inf [10-Inf] | $3.3 \times 10^{-7}$ |
| rs113318830 | 2 | 11897940 | *LPIN1* | - | AT | 0.78 | 1.25 [1.14-1.36] | $3.7 \times 10^{-7}$ |
| rs796733043 | 7 | 125639265 | *GRM8* | TATT | - | 0.82 | 1.32 [1.18-1.47] | $7.7 \times 10^{-7}$ |
| rs142956186 | 4 | 163162586 | *FSTL5* | T | C | $1.7 \times 10^{-3}$ | Inf [11.17-Inf] | $9.9 \times 10^{-7}$ |

Loci with $p < 1 \times 10^{-6}$ with endometriosis-related infertility
Chr: chromosome, Pos: position, RA: risk allele, OA: Other allele, RAF: Risk allele frequency, OR: Odds ratio, P: P-value

**Figure 2.**
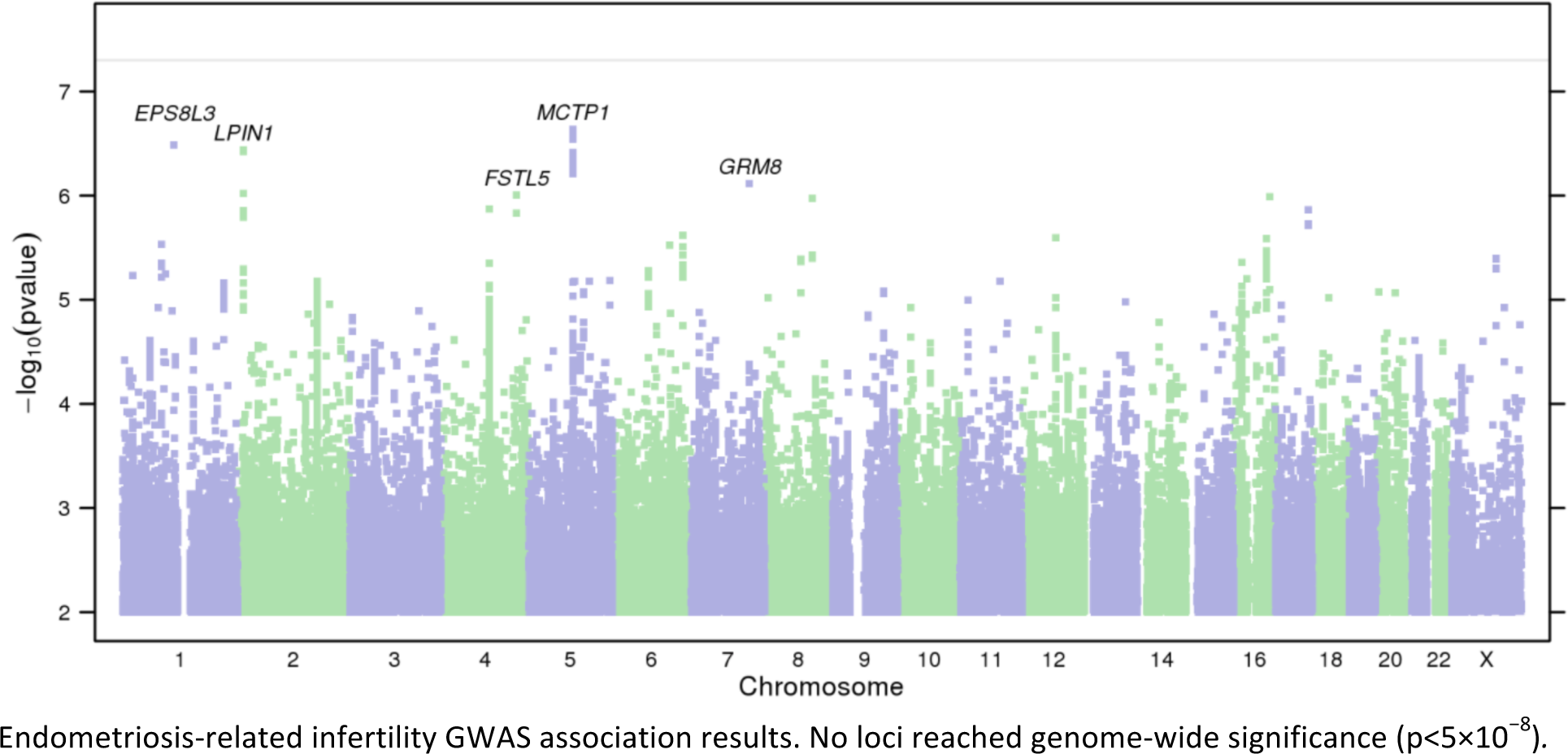
Manhattan plot of endometriosis-related infertility GWAS.

## Discussion

### Self-reported endometriosis phenotype

Although the endometriosis phenotype in our study was self-reported, seven endometriosis loci previously identified in GWAS were replicated at genome-wide significance and four were replicated at a nominal level with a consistent direction of effect. Only two loci did not replicate (Tables 3-4). These results are concordant with the reported accuracy of self-reported endometriosis phenotypes in comparison with the Swedish National Inpatient Registry with a specificity of 97.0%, a sensitivity of 61.8% and receiver operating characteristics area of 0.79 [27]. However, replicating our novel loci in datasets with surgically confirmed cases is important to avoid identification of loci associated with dysmenorrhea in the absence of endometriosis. Interestingly, 22.2% of women with endometriosis-related infertility reported having been diagnosed with and/or treated for polycystic ovary syndrome (PCOS), compared to 14.8% of fertile women with endometriosis (Table 2). Because of the complexity of the PCOS diagnosis and the self-reported nature of the PCOS phenotype in our dataset, it is possible that some of the women who reported having PCOS, had anovulation due to their endometriosis [28], which could contribute to their infertility. It is also possible that PCOS contributes to the infertility of some of our endometriosis-related infertility cases.

### Novel endometriosis loci

Seven novel loci were associated with endometriosis at genome-wide significance in our analysis. The novel locus *NGF* was also associated at genome-wide significance with severity of dysmenorrhea in another study of 23andMe female participants of European ancestry [29]. Interestingly, the expression level of *NGF* in the peritoneal fluid has been observed to be higher in women with endometriosis, and blocking NGF significantly decreases neurite outgrowth in endometriotic lesions [30]. Multiple transcription factors have been found to bind the region covered by the credible SNPs set within the *NGF* locus (Table 6). This region also overlaps with predicted enhancers in multiple tissues, including the fetal adrenal gland, fetal stomach, fetal kidney, colon smooth muscle, adipose nuclei, fetal lung, primary monocytes from peripheral blood, fetal muscle trunk, and fetal muscle leg.

**Table 6.** Transcription factors binding the DNA region covered by the credible set according to ChiP-seq in the Encode data

| Locus | Transcription factors |
| --- | --- |
| *NGF* | AR, FOXP2, RFX3, SPI1 |
| *ATP1B1-F5* | RFX3 |
| CD109 | CTCF, SETDB1, E2F4, FOS, RAD21, SMC3, CEBPB, EP300, JUND, MAFK, RFX5, TEAD4, EGR1, NFYB |
| *HEY2* | BATF, GATA3, FOXA1, POLR2A, RFX3, CEBPB, TRIM28, CTCF, RAD21, EBF1, ATF2, TAF7, GATA2, FOS, MAFF, MAFK, CHD1, GTF2F1, SETDB1 |
| *OSR2-VPS13B* | ARID3A, ATF1, ATF2, ATF3, BATF, BCL11A, CBX3, CCNT2, CHD1, CTPBP2, EGR1, ESR1, EZH2, FOS, FOSL1, FOXA1, FOXA2, CEBPB, EP300, FOS, FOXM1, GABPB1, GATA1, GATA2, GATA3, HDAC2, IRF1, IRF4, JUN, JUNB, JUND, MAFF, MAFK, MAX, MTA3, MYC, NFKB1, NFIC, NR2F2, POLR2A, POU5F1, RCOR1, REST, RUNX3, SETDB1, SPDEF, SPI1, STAT5A, SUZ12, TAF1, TAL1, TBL1XR1, TCF3, TCF7L2, TCF12, TEAD4, TRIM28, ZNF143, ZNF217, ZNF263 |
| *WT1* | AR, ARID3A, ATF3, CBX3, CEBPB, CTCF, E2F6, EP300, ESR1, FOS, FOXA1, GTF2F1, IKZF1, JUND, MAFF, MAFK, MAX, MXI1, MYC, NR3C1, POLR2A, RAD21, RFX5, SIN3A, SMARCC1, SMARCC2, SMC3, TAF1, USF1, USF2, ZNF143, ZNF263 |
| *TEX11-SLC7A3* | CEBPB, CTCF, ELF1, EP300, GABPB1, MXI1, NR2F2, PML, RCOR1, STAT5A, TAL1, TEAD4, USF1, YY1 |

The credible SNPs set of the second novel endometriosis locus, *ATP1B1-F5*, includes the Factor V Leiden (*F5*) p.Gln534Arg missense variant rs6025, which has been associated with venous thromboembolism [31-33], thrombosis [34], inflammatory bowel disease [35], and recurrent pregnancy loss [36], although the risk allele for endometriosis is inverted.

The third novel endometriosis locus is near the gene *CD109*, which encodes a glycosyl phosphatidylinositol (GPI)-linked glycoprotein that negatively regulates signaling by transforming growth factor beta (TGFB1). *TGFB1* is thought to be involved in different aspects of the pathophysiology of endometriosis including endometrial cell proliferation and tissue remodeling [37,38].

The fourth novel endometriosis locus is near the gene Hes Related Family bHLH Transcription Factor with YRPW Motif 2 (*HEY2*), the expression of which is downregulated by 17β-estradiol in ovariectomized mice [39]. The nearby gene *NCOA7* modulates the activity of the estrogen receptor (ER) [40]. The locus has been associated at genome-wide significance with endometrial [41] and breast [42] cancer. It is an eQTL for both *HEY2* and *NCOA7*, is bound by multiple transcription factors (Table 6), and contains predicted enhancers in multiple tissues including the ovary, psoas muscle, aorta, left ventricle, skeletal muscle in females, fetal lung, fetal adrenal gland, and fetal heart.

The fifth novel endometriosis locus is *OSR2-VPS13B*. The gene odd-skipped related transcription factor 2 (*OSR2*) is a transcription factor that is highly expressed in the endometrium and the ovary, and its expression is down-regulated upon progesterone receptor knockdown in human endometrial stromal cells, suggesting that it may be involved in decidualization [43]. In a genome-wide epigenetics study based on CpG methylation, *OSR2* showed increased methylation and decreased expression in endometriotic stromal cells compared to stromal cells from normal endometrium [44]. Over 40 transcription factors have been identified as binding the region covered by the credible SNP set (Table 6), and the region also overlaps with predicted enhancers in numerous tissues. The variant rs3019295, part of the credible SNP set, is in an activated transcription site of *OSR2* in the ovary, psoas muscle, and aorta.

The sixth novel endometriosis locus is near the gene Wilms tumor 1 (*WT1*). *WT1* encodes a transcription factor that is highly expressed in the endometrium, the testis, and the ovary, and that has an essential role in the normal development of the urogenital system. It is selectively expressed in neurons of deep endometriosis [45] and is down-regulated in endometriotic stromal cells compared to endometrial stromal cells [46,47]. The region covered by the credible SNP set overlaps with the binding sites of several transcription factors and with predicted enhancers in multiple tissues.

The seventh novel endometriosis locus is *TEX11*-*SLC7A3*. The gene solute carrier family 7 member 3 (*SLC7A3*) encodes a sodium-independent cationic amino acid transporter, and it is highly expressed in the endometrium. Testis expressed 11 (*TEX11*) is expressed in the pancreas, testis, and human fetal ovary, and is associated with azoospermia and male infertility [48-52]. TEX11 competes with estrogen receptor beta (ERβ) for binding to hematopoietic pre-B cell leukemia transcription factor-interacting protein (HPIP), which modulates the function of ERs [53]. TEX11 suppresses estrogen-stimulated germ cell proliferation and affects the expression of estrogen target genes through its binding to HPIP [53].

### Endometriosis-related infertility association results

No loci reached genome-wide significance in the endometriosis-related infertility GWAS. Of the 14 genome-wide endometriosis loci, only four showed nominal association with endometriosis-related infertility, suggesting that the genetic factors involved in endometriosis and endometriosis-related infertility may differ.

The most strongly associated locus with endometriosis-related infertility is a rare (minor allele frequency of 8.8×10^-3^) insertion in the intron of the gene Multiple C2 and Transmembrane Domain Containing 1 (*MCTP1*). The homolog of *MCTP1* in *Caenorhabditis elegans* (D2092.1) is an essential gene, and its ablation leads to early embryonic lethality [54]. The region covered by the credible set is predicted to contain an enhancer in human umbilical vein endothelial primary cells, fetal lung, fetal muscle leg, and fetal stomach. The protein encoded by *MCTP1* is not well characterized, but it is an evolutionarily conserved protein that contains C_2_ domains of high Ca^2+^-binding affinity. It may have a Ca^2+^-controlled regulatory function or serve as a Ca^2+^ buffer [55]. Functional studies in central nervous system neurons showed that *MCTP1* over-expression significantly inhibited neuronal transferrin endocytosis, secretory vesicle retrieval, cell migration, and oxidative stress from glutamate toxicity [56]. If confirmed, the association of this locus with endometriosis-related infertility might involve *MCTP1* protecting oocyte homeostasis, maturation, and fertilization from the oxidative stress generated by endometriosis.

The second most strongly associated locus in the endometriosis-related infertility GWAS was *EPS8L3-CSF1.* The gene *CSF1* encodes an estrogen-regulated cytokine that controls the production, differentiation, and function of macrophages. In humans, CSF1 level is increased in the pregnant endometrium compared to the nonpregnant and is high in the placenta throughout pregnancy [57,58]. In mice, females lacking CSF1 have extended estrus cycles and poor ovulation rates [59,60]. CSF1 levels are increased in the peritoneal fluid of patients with endometriosis [61] and *Csf1* expression was significantly higher in lesions of an endometriosis mouse model [62]. The ER-dependent regulation of *CSF1* in peripheral nerve fibers has been suggested to play a critical role in early development of endometriotic lesions [63] and in modulating macrophage survival [62]. Given the potential role of oxidative stress in endometriosis-related infertility, it is also worth noting that the oxidative stress response genes *GSTM1, GSTM2, GSTM3, GSTM4*, and *GSTM5* are < 200kb from the associated variant in this locus, although a high recombination rate separates the genes from the top variant.

The third most strongly associated locus in the endometriosis-related infertility GWAS was *LPIN1*, which acts as a proinflammatory mediator during TLR signaling and during the development of in vivo inflammatory processes [64]. In mice, *Lpin1* is down-regulated in the uterus by estradiol via the ER [65] and *Lpin1* deficiency is associated with impaired fertility [66,67]. Lipin1 might also play a role in the regulation of the uterine cell cycle because it has been found to have anti-proliferative effects in murine pro B cells [68].

## Conclusion

Our study replicated seven loci identified in previous endometriosis GWASs and identified seven new candidate loci with genome-wide significance. These new loci need to be replicated, and further studies are needed to determine the causal variants. Although no loci had genome-wide significance for endometriosis-related infertility in this study, the three most strongly associated loci were in or near genes with roles in female fertility or embryonic lethality in model organisms. These genes are thought to be involved in oxidative stress, macrophage survival, and inflammation. These results provide new candidate genes involved in the pathophysiology of endometriosis and endometriosis-related infertility.

## Acknowledgements

We thank the research participants and employees of Celmatix and 23andMe who contributed to this study. We acknowledge the contributions from additional members of the Celmatix Research Team, including R. Mark Adams, Daniela S. Colaci, Chris Glazner, Samuel T. Globus, Sean O’Keeffe, Ursula M. Schick, Lei Tan, Cameron D. Wellock, Danielle White, and Rajeshwari R. Valiathan. Members of the 23andMe Research Team are: Michelle Agee, Babak Alipanahi, Adam Auton, Robert K. Bell, Katarzyna Bryc, Sarah L. Elson, Nicholas A. Furlotte, David A. Hinds, Karen E. Huber, Aaron Kleinman, Nadia K. Litterman, Matthew H. McIntyre, Joanna L. Mountain, Elizabeth S. Noblin, Carrie A.M. Northover, Steven J. Pitts, J. Fah Sathirapongsasuti, Olga V. Sazonova, Janie F. Shelton, Suyash Shringarpure, Chao Tian, Vladimir Vacic, Catherine H. Wilson.

## Data availability

The full GWAS summary statistics for the 23andMe data sets will be made available through 23andMe to qualified researchers under an agreement with 23andMe that protects the privacy of the 23andMe participants. Please visit research.23andMe.com/collaborate for more information and to apply to access the data.

## References

1. Jenkins, S., D.L. Olive, and A.F. Haney, Endometriosis: pathogenetic implications of the anatomic distribution. Obstet Gynecol, 1986. 67(3): p. 335-8.

2. Gao, X., et al., Economic burden of endometriosis. Fertil Steril, 2006. 86(6): p. 1561–72.

3. Bulletti, C., et al., Endometriosis and infertility. J Assist Reprod Genet, 2010. 27(8): p. 441–7.

4. Barnhart, K., R. Dunsmoor-Su, and C. Coutifaris, Effect of endometriosis on in vitro fertilization. Fertil Steril, 2002. 77(6): p. 1148–55.

5. Akande, V.A., et al., Differences in time to natural conception between women with unexplained infertility and infertile women with minor endometriosis. Hum Reprod, 2004. 19(1): p. 96–103.

6. Johnson, N.P., et al., The FLUSH trial--flushing with lipiodol for unexplained (and endometriosis-related) subfertility by hysterosalpingography: a randomized trial. Hum Reprod, 2004. 19(9): p. 2043–51.

7. Jansen, R.P., Minimal endometriosis and reduced fecundability: prospective evidence from an artificial insemination by donor program. Fertil Steril, 1986. 46(1): p. 141–3.

8. Simon, C., et al., Outcome of patients with endometriosis in assisted reproduction: results from in-vitro fertilization and oocyte donation. Hum Reprod, 1994. 9(4): p. 725–9.

9. Khine, Y.M., F. Taniguchi, and T. Harada, Clinical management of endometriosis-associated infertility. Reprod Med Biol, 2016. 15(4): p. 217–225.

10. Gajbhiye, R., et al., Multiple endometrial antigens are targeted in autoimmune endometriosis. Reprod Biomed Online, 2008. 16(6): p. 817–24.

11. Osborn, B.H., et al., Inducible nitric oxide synthase expression by peritoneal macrophages in endometriosis-associated infertility. Fertil Steril, 2002. 77(1): p. 46–51.

12. Agarwal, A., et al., The effects of oxidative stress on female reproduction: a review. Reprod Biol Endocrinol, 2012. 10: p. 49.

13. Marcoux, S., R. Maheux, and S. Berube, Laparoscopic surgery in infertile women with minimal or mild endometriosis. Canadian Collaborative Group on Endometriosis. N Engl J Med, 1997. 337(4): p. 217–22.

14. Treloar, S.A., et al., Genetic influences on endometriosis in an Australian twin sample. Fertil Steril, 1999. 71(4): p. 701–10.

15. Saha, R., et al., Heritability of endometriosis. Fertil Steril, 2015. 104(4): p. 947–952.

16. Adachi, S., et al., Meta-analysis of genome-wide association scans for genetic susceptibility to endometriosis in Japanese population. J Hum Genet, 2010. 55(12): p. 816–21.

17. Painter, J.N., et al., Genome-wide association study identifies a locus at 7p15.2 associated with endometriosis. Nat Genet, 2011. 43(1): p. 51–4.

18. Nyholt, D.R., et al., Genome-wide association meta-analysis identifies new endometriosis risk loci. Nat Genet, 2012. 44(12): p. 1355–9.

19. Albertsen, H.M., et al., Genome-wide association study link novel loci to endometriosis. PLoS One, 2013. 8(3): p. e58257.

20. Rahmioglu, N., et al., Genetic variants underlying risk of endometriosis: insights from meta-analysis of eight genome-wide association and replication datasets. Hum Reprod Update, 2014. 20(5): p. 702–16.

21. Steinthorsdottir, V., et al., Common variants upstream of KDR encoding VEGFR2 and in TTC39B associate with endometriosis. Nat Commun, 2016. 7: p. 12350.

22. Sapkota, Y., et al., Meta-analysis identifies five novel loci associated with endometriosis highlighting key genes involved in hormone metabolism. Nat Commun, 2017. 8: p. 15539.

23. Uno, S., et al., A genome-wide association study identifies genetic variants in the CDKN2BAS locus associated with endometriosis in Japanese. Nat Genet, 2010. 42(8): p. 707–10.

24. Wakefield, J., A Bayesian measure of the probability of false discovery in genetic epidemiology studies. Am J Hum Genet, 2007. 81(2): p. 208–27.

25. Wellcome Trust Case Control, C., et al., Bayesian refinement of association signals for 14 loci in 3 common diseases. Nat Genet, 2012. 44(12): p. 1294–301.

26. Machiela, M.J. and S.J. Chanock, LDlink: a web-based application for exploring population-specific haplotype structure and linking correlated alleles of possible functional variants. Bioinformatics, 2015. 31(21): p. 3555–7.

27. Saha, R., L. Marions, and P. Tornvall, Validity of self-reported endometriosis and endometriosis-related questions in a Swedish female twin cohort. Fertil Steril, 2017. 107(1): p. 174–178 e2.

28. Soules, M.R., et al., Endometriosis and anovulation: a coexisting problem in the infertile female. Am J Obstet Gynecol, 1976. 125(3): p. 412–7.

29. Jones, A.V., et al., Genome-wide association analysis of pain severity in dysmenorrhea identifies association at chromosome 1p13.2, near the nerve growth factor locus. Pain, 2016. 157(11): p. 2571–2581.

30. Barcena de Arellano, M.L., et al., Overexpression of nerve growth factor in peritoneal fluid from women with endometriosis may promote neurite outgrowth in endometriotic lesions. Fertil Steril, 2011. 95(3): p. 1123–6.

31. Heit, J.A., et al., A genome-wide association study of venous thromboembolism identifies risk variants in chromosomes 1q24.2 and 9q. J Thromb Haemost, 2012. 10(8): p. 1521– 31.

32. Germain, M., et al., Meta-analysis of 65,734 individuals identifies TSPAN15 and SLC44A2 as two susceptibility loci for venous thromboembolism. Am J Hum Genet, 2015. 96(4): p. 532–42.

33. Klarin, D., et al., Genetic Analysis of Venous Thromboembolism in UK Biobank Identifies the ZFPM2 Locus and Implicates Obesity as a Causal Risk Factor. Circ Cardiovasc Genet, 2017. 10(2).

34. Hinds, D.A., et al., Genome-wide association analysis of self-reported events in 6135 individuals and 252 827 controls identifies 8 loci associated with thrombosis. Hum Mol Genet, 2016. 25(9): p. 1867–74.

35. Liu, J.Z., et al., Association analyses identify 38 susceptibility loci for inflammatory bowel disease and highlight shared genetic risk across populations. Nat Genet, 2015. 47(9): p. 979–986.

36. Sergi, C., T. Al Jishi, and M. Walker, Factor V Leiden mutation in women with early recurrent pregnancy loss: a meta-analysis and systematic review of the causal association. Arch Gynecol Obstet, 2015. 291(3): p. 671–9.

37. Dela Cruz, C. and F.M. Reis, The role of TGFbeta superfamily members in the pathophysiology of endometriosis. Gynecol Endocrinol, 2015. 31(7): p. 511–5.

38. Young, V.J., et al., The role of TGF-beta in the pathophysiology of peritoneal endometriosis. Hum Reprod Update, 2017. 23(5): p. 548–559.

39. Nakamura, T., et al., Sequential changes in the expression of Wnt-and Notch-related genes in the vagina and uterus of ovariectomized mice after estrogen exposure. In Vivo, 2012. 26(6): p. 899–906.

40. Shao, W., S. Halachmi, and M. Brown, ERAP140, a conserved tissue-specific nuclear receptor coactivator. Mol Cell Biol, 2002. 22(10): p. 3358–72.

41. Cheng, T.H., et al., Five endometrial cancer risk loci identified through genome-wide association analysis. Nat Genet, 2016. 48(6): p. 667–74.

42. Higginbotham, K.S., et al., A multistage association study identifies a breast cancer genetic locus at NCOA7. Cancer Res, 2011. 71(11): p. 3881–8.

43. Cloke, B., et al., The androgen and progesterone receptors regulate distinct gene networks and cellular functions in decidualizing endometrium. Endocrinology, 2008. 149(9): p. 4462–74.

44. Yotova, I., et al., Epigenetic Alterations Affecting Transcription Factors and Signaling Pathways in Stromal Cells of Endometriosis. PLoS One, 2017. 12(1): p. e0170859.

45. Coosemans, A., et al., Wilms’ tumor gene 1 (WT1) overexpression in neurons in deep endometriosis: a pilot study. Fertil Steril, 2009. 91(4 Suppl): p. 1441-4.

46. Gurates, B., et al., WT1 and DAX-1 inhibit aromatase P450 expression in human endometrial and endometriotic stromal cells. J Clin Endocrinol Metab, 2002. 87(9): p. 4369–77.

47. Matsuzaki, S., et al., Expression of WT1 is down-regulated in eutopic endometrium obtained during the midsecretory phase from patients with endometriosis. Fertil Steril, 2006. 86(3): p. 554–8.

48. Sha, Y., et al., A novel TEX11 mutation induces azoospermia: a case report of infertile brothers and literature review. BMC Med Genet, 2018. 19(1): p. 63.

49. Yang, F., et al., TEX11 is mutated in infertile men with azoospermia and regulates genome-wide recombination rates in mouse. EMBO Mol Med, 2015. 7(9): p. 1198–210.

50. Yatsenko, A.N., et al., X-linked TEX11 mutations, meiotic arrest, and azoospermia in infertile men. N Engl J Med, 2015. 372(22): p. 2097–107.

51. Nakamura, S., et al., Next-generation sequencing for patients with non-obstructive azoospermia: implications for significant roles of monogenic/oligogenic mutations. Andrology, 2017. 5(4): p. 824–831.

52. Zhang, X., et al., Six polymorphisms in genes involved in DNA double-strand break repair and chromosome synapsis: association with male infertility. Syst Biol Reprod Med, 2015. 61(4): p. 187–93.

53. Yu, Y.H., et al., TEX11 modulates germ cell proliferation by competing with estrogen receptor beta for the binding to HPIP. Mol Endocrinol, 2012. 26(4): p. 630–42.

54. Maeda, I., et al., Large-scale analysis of gene function in Caenorhabditis elegans by high-throughput RNAi. Curr Biol, 2001. 11(3): p. 171–6.

55. Shin, O.H., et al., Evolutionarily conserved multiple C2 domain proteins with two transmembrane regions (MCTPs) and unusual Ca2+ binding properties. J Biol Chem, 2005. 280(2): p. 1641–51.

56. Qiu, L., H. Yu, and F. Liang, Multiple C2 domains transmembrane protein 1 is expressed in CNS neurons and possibly regulates cellular vesicle retrieval and oxidative stress. J Neurochem, 2015. 135(3): p. 492–507.

57. Daiter, E., et al., Expression of colony-stimulating factor-1 in the human uterus and placenta. J Clin Endocrinol Metab, 1992. 74(4): p. 850–8.

58. Kauma, S.W., et al., Colony-stimulating factor-1 and c-fms expression in human endometrial tissues and placenta during the menstrual cycle and early pregnancy. J Clin Endocrinol Metab, 1991. 73(4): p. 746–51.

59. Cohen, P.E., et al., Colony-stimulating factor 1 regulation of neuroendocrine pathways that control gonadal function in mice. Endocrinology, 2002. 143(4): p. 1413–22.

60. Ovadia, S., K. Insogna, and G.Q. Yao, The cell-surface isoform of colony stimulating factor 1 (CSF1) restores but does not completely normalize fecundity in CSF1-deficient mice. Biol Reprod, 2006. 74(2): p. 331–6.

61. Fukaya, T., et al., The role of macrophage colony stimulating factor in the peritoneal fluid in infertile patients with endometriosis. Tohoku J Exp Med, 1994. 172(3): p. 221–6.

62. Greaves, E., et al., Estradiol is a critical mediator of macrophage-nerve cross talk in peritoneal endometriosis. Am J Pathol, 2015. 185(8): p. 2286–97.

63. Jensen, J.R., et al., A potential role for colony-stimulating factor 1 in the genesis of the early endometriotic lesion. Fertil Steril, 2010. 93(1): p. 251–6.

64. Meana, C., et al., Lipin-1 integrates lipid synthesis with proinflammatory responses during TLR activation in macrophages. J Immunol, 2014. 193(9): p. 4614–22.

65. Gowri, P.M., et al., Lipin1 regulation by estrogen in uterus and liver: implications for diabetes and fertility. Endocrinology, 2007. 148(8): p. 3685–93.

66. Reue, K. and M.H. Doolittle, Naturally occurring mutations in mice affecting lipid transport and metabolism. J Lipid Res, 1996. 37(7): p. 1387–405.

67. Peterfy, M., et al., Lipodystrophy in the fld mouse results from mutation of a new gene encoding a nuclear protein, lipin. Nat Genet, 2001. 27(1): p. 121–4.

68. Brachat, A., et al., A microarray-based, integrated approach to identify novel regulators of cancer drug response and apoptosis. Oncogene, 2002. 21(54): p. 8361–71.

